# Siderophore-mediated inhibition of *Legionella pneumophila* by environmental *Pseudomonas* isolates

**DOI:** 10.64898/2026.01.19.700273

**Authors:** Alessio Cavallaro, Johanna Kohler, Marco Gabrielli, Vera Vollenweider, Rolf Kümmerli, Frederik Hammes

**Author notes:** Corresponding author: Name: Frederik Hammes. These authors contributed equally (Alessio Cavallaro, Johanna Kohler).

## Abstract

The genus *Legionella* comprises opportunistic pathogenic bacteria commonly found in natural and engineered water systems, where they interact with environmental microbes and protozoa, primarily in biofilms. *Legionella pneumophila* is the main causative agent of Legionnaires’ disease and is transmitted through inhalation of contaminated aerosols. Iron availability is a critical factor for *L. pneumophila* growth, persistence, and virulence, yet iron is often limited in aquatic environments. To overcome iron scarcity, many bacteria produce siderophores, secondary metabolites that scavenge ferric iron. Because siderophores are chemically diverse and species specific, they play a key role in inter-species competition and can withhold iron from competitors.

Here, we investigated the effects of iron depletion and siderophore-mediated competition on *L. pneumophila* using commercial pyoverdines and extracellular metabolites from environmental *Pseudomonas* strains. Growth assays showed that *L. pneumophila* can grow under iron-limited conditions but with lag phases extended by more than 20 hours. Pyoverdines inhibited growth in a concentration-dependent manner, primarily increasing the time to mid-log phase (t_mid). Supernatants and crude pyoverdine extracts from siderophore-producing *Pseudomonas* strains caused the strongest inhibition, including lag-phase extensions of up to 55 hours or complete growth arrest. These results demonstrate that siderophore-producing bacteria can suppress *L. pneumophila* by limiting iron availability.

## 1. Introduction

Bacteria belonging to the genus *Legionella* are opportunistic pathogens ubiquitous in freshwater environments (Fields *et al*., 2002, Mondino *et al*., 2020). *Legionella pneumophila* is the most virulent species, associated with > 92% of confirmed clinical cases (Hammes *et al*., 2025). Human infection occurs through inhalation of *Legionella*-containing aerosols, leading to either Legionnaires’ disease, a severe pneumonia with a high case fatality rate in vulnerable populations or Pontiac fever, a mild-cold with flu-like symptoms (Fields *et al*., 2002). *Legionella* often colonize engineered water systems like cooling towers and building plumbing systems, that ultimately represent the main exposure route for humans (Falkinham *et al*., 2015). In such engineered aquatic systems, *Legionella* are embedded in biofilms (Declerck, 2010, Barbosa *et al*., 2024), exhibiting a biphasic lifestyle that includes replication within a host cell such as Amoeba or other protists (Boamah *et al*., 2017, Mondino *et al*., 2020) and survival in biofilms as free-living cells.

Iron availability is believed to be a major factor in the replication and survival of *L. pneumophila* (Gebran *et al*., 1994, Cianciotto, 2007). It has been demonstrated that host-derived iron contributes to bacterial growth (Isaac *et al*., 2015), while iron limitation attenuates virulence gene expression, restricts bacterial replication, and can even trigger premature intracellular escape from the host cell (O’Connor Tamara *et al*., 2016). *L. pneumophila* possesses different iron uptake systems, including the ferrous transporter FeoB (Robey & Cianciotto, 2002) and the ferric iron siderophore legiobactin (Liles *et al*., 2020), whose biosynthesis and transport are regulated under low-iron conditions. Additionally, it has been showed that iron modulates the expression of at least one effector belonging to the Dot/Icm(Vollenweider *et al*., 2024) secretion system, a type IV secretion system required for the establishment inside a host and for starting the replicative phase (Isaac *et al*., 2015). This process is mediated by the ferric uptake regulator (Fur), and creates an important link between iron metabolism and virulence regulation (Portier *et al*., 2015)

When the bioavailability of iron is low, for instance, when it occurs in insoluble or host-bound forms, many bacterial genera produce siderophores to scavenge and solubilize ferric iron, a mechanism well described for a wide range of bacteria, as well as fungi and some plants (Hider & Kong, 2010). A comprehensive database of 649 unique siderophores structures has been developed by He and colleagues (He *et al*., 2024). *Pseudomonas* isolates are well-document for using siderophores to scavenge iron from the environment (Cornelis, 2010). Siderophores are iron-chelating molecules secreted by bacteria to sequester ferric iron (Schalk, 2025). Once the iron is bound to the siderophores, it is acquired by the bacteria through membrane receptors, which are specific for each type of siderophores (Schalk, 2025). Their secretion into the environment creates opportunities for different types of interaction among strains and species (Figueiredo *et al*., 2022). Since closely related strains or species typically produce the same siderophore (e.g. enterobactin produced by several Enterobacteriales species), the secreted molecules can be shared within the community and used as public good by everyone with compatible receptors. Conversely, different species typically produce structurally distinct siderophores with incompatible receptor systems. In such cases, species use siderophores for competition in iron-limited niches (Niehus *et al*., 2017). Certain siderophores are especially potent to withhold iron from competitors due to the exceptionally high stability of their ferric complexes (Vollenweider *et al*., 2024). Because of their competitive nature in inter-species interactions, siderophores have recently been proposed as tools to restrict the growth of unwanted bacteria. For example, several pyoverdines, siderophores isolated from benign environmental *Pseudomonas* spp., have been shown to suppress human pathogens such as *Acinetobacter baumannii* and *Staphylococcus aureus* by depriving them of iron (Vollenweider *et al*., 2024).

In the present study, we aim to extend this approach to the inhibition of *Legionella* spp.. Although the iron dependency of many *Legionella* species is well established, the specific effects of iron availability on its growth, as well as its response to iron competition mediated by siderophores, remain poorly understood. Here, we address this gap by directly testing the impact of iron depletion and siderophore activity on *L. pneumophila*. We first tested the growth of *L. pneumophila* and other *Legionella* species under variable iron conditions, identifying the effects of iron starvation on *Legionella* growth. Subsequently, we used a commercially available pyoverdine and crude-purified pyoverdines from extracellular extracts of environmental *Pseudomonas* strains to quantify the inhibition of *L. pneumophila* in under iron depleted or repleted conditions.

## 2.2 Material and Methods

### 2.1. Bacterial strains

#### 2.1.1 *Legionella* strains

Five *Legionella* species (*L. pneumophila* (serogroup 1 (Lp1)*), L*. *londiniensis, L. jordanis, L. longbeachae, L. feeleii)* were used to assess growth responses to iron availability and susceptibility to siderophore-mediated inhibition. All species were obtained from the National *Legionella* Reference Center (CNRL) located in Bellinzona, Switzerland.

#### 2.1.2 Siderophore-producing *Pseudomonas* strains

Environmental *Pseudomonas* isolates were originally obtained from 16 independent water and soil samples collected in Campus Irchel Park (University of Zurich). All isolates are non-pathogenic and were previously described (Butaitė *et al*., 2017, Butaitė *et al*., 2018, Kramer *et al*., 2019). The five strains selected for this study were chosen based on their production of potent pyoverdines, as confirmed by a two-step functional screen detailed in Vollenweider et al. (Vollenweider *et al*., 2024). Briefly, culture supernatants from 320 *Pseudomonas* isolates were screened for their ability to inhibit 12 human pathogens. To identify siderophore-mediated effects, 40 μM FeCl₃ was added to the supernatants in a second screening — pyoverdine-dependent inhibition was expected to diminish under these iron-replete conditions. This approach identified seven strong candidates producing five chemically distinct pyoverdines. For the present study, one representative strain per pyoverdine type was selected, namely *Pseudomonas s3b09*, *3G07, 3A06, s3c13,* and *s3e20 (Vollenweider et al., 2024)* .

### 2.2. Preparation of *Pseudomonas* supernatants

Five *Pseudomonas* strains (s3b09, 3G07, 3A06, s3c13, s3e20) were grown overnight in 3 mL of LB broth at 28 °C with orbital shaking at 170 rpm (triplicate cultures per strain). Following incubation, 250 μL from each replicate was transferred into 25 mL of iron-depleted medium composed of 1% casamino acids (CAA) supplemented with 250 μM 2,2’-bipyridyl (Sigma-Aldrich) in 100 mL Erlenmeyer flasks. Cultures were incubated overnight at 28 °C, 170 rpm. After growth, siderophore production was confirmed by measuring both optical density at 600 nm (OD₆₀₀) and fluorescence (excitation 400 nm, emission 460 nm) using a microplate reader (96-well format, 200 μL per replicate). To obtain cell-free supernatants, cultures were first centrifuged at 2250 rcf for 10 minutes (Eppendorf 5910 Ri, Germany). Supernatants were transferred to 50 mL Greiner tubes and centrifuged again at 4300 rcf for 10 minutes to ensure removal of residual cells. The resulting supernatants were sterile-filtered (0.2 μm,) Sarstedt, Germany), aliquoted (2 mL per tube), and stored at –20 °C until use in downstream 96-well growth assays.

### 2.3. Growth assays in 96-well plates

Prior to experimentation, *Legionella* strains were pre-cultured on buffered charcoal yeast extract (BCYE) agar for 72 hours at 37 °C with orbital shaking. Colonies were then transferred to liquid media for culture preparation. Following incubation for 72 hours, *Legionella* cultures were centrifuged at 2500 rcf for 5 minutes (Hettich Rotina 380, Switzerland), washed twice in phosphate-buffered saline (PBS), and resuspended to an OD₆₀₀ of 0.1 (Biochrom Libra S4, UK). For inoculation, cultures were further diluted to a final OD₆₀₀ of 0.01 in experimental wells.

Growth assays were conducted in 96-well Nunc Edge 2.0 plates (Thermo Scientific, USA) using a BioTek Synergy H1 plate reader (Agilent, USA). Each well contained 180 μL of medium with the respective treatment (e.g., iron chelator, commercial pyoverdines (Sigma-Aldrich), *Pseudomonas* supernatant or extract), and was inoculated with 20 μL of pre-diluted *Legionella* suspension.

Plates were incubated at 37 °C with continuous linear shaking at 567 cpm. Optical density at 600 nm was measured every 6 minutes for up to 99 hours. To minimize evaporation, the plate’s peripheral moats were filled with 1.7 mL sterile water. For each treatment condition, triplicate wells were included. Negative controls consisted of uninoculated wells containing PBS in place of *Legionella* suspension.

### 2.4. Supernatant inhibition assay

To assess the inhibitory effects of extracellular products from *Pseudomonas* isolates, cell-free supernatants obtained under iron-limited conditions were tested in a liquid growth assay. For each treatment, 18□μL of *Pseudomonas* supernatant was added to 162□μL of iron-supplemented BYEB medium (BYEB□+□Fe) in a 96-well plate. Two controls were included: 1) Medium-only control: 180□μL BYEB□+□Fe without additives. 2) PBS control: 18□μL PBS added to 162□μL BYEB□+□Fe, simulating nutrient dilution introduced by spent medium. Each condition was inoculated with 20 μL of *L. pneumophila*, and growth was monitored under the conditions described in Section 2.3. To evaluate whether the observed inhibition was primarily mediated by iron sequestration via pyoverdines, supernatants were also added to unsupplemented BYEB (iron-limited) in parallel assays.

### 2.5. Crude pyoverdine inhibition assay

Crude pyoverdine extracts were prepared as described in Vollenweider et al. (Vollenweider *et al*., 2024). Briefly, supernatants from the five *Pseudomonas* strains were processed using Amberlite XAD-16N resin (Sigma-Aldrich, Switzerland) columns. Fractions with the highest fluorescence (excitation: 400□nm, emission: 460□nm) — indicating peak pyoverdine content — were pooled, and the methanolic eluent was evaporated. Samples were lyophilized and stored at –20□°C. As the extracts were not chemically purified, concentrations were expressed in mg/mL. Lyophilized powders were resuspended in BYEB□+□Fe, sterile filtered (0.2□μm syringe filters), and adjusted to a 10□mg/mL stock solution. Working dilutions were prepared in BYEB□+□Fe or BYEB to final concentrations of 1□mg/mL and 0.5□mg/mL, as appropriate. Growth inhibition assays were performed in 96-well plates by adding the diluted pyoverdine preparations to BYEB□+□Fe or BYEB, then inoculating with *L. pneumophila* as described in Section 2.3. Each condition was tested in duplicate plates with appropriate untreated controls on each plate.

### 2.6. Data analysis

Growth curves obtained from OD₆₀₀ measurements were analysed using RStudio Version 4.4.1 (RStudio Team, 2024). Growth metrics were calculated using the growthcurver R package (Version 0.3.1 (Sprouffske & Wagner, 2016)), which fits a logistic growth model to microbial time-course data. The primary metrics extracted were: 1) Intrinsic growth rate (*r*); 2) Carrying capacity (*K*); 3) *t*_mid: the time at which OD₆₀₀ reaches half of the maximum observed value (used as a proxy for lag phase duration). Before analysis, background absorbance was corrected by subtracting the time-matched mean OD₆₀₀ of corresponding control wells from each treatment well. To assess differences in growth parameters across conditions (e.g., varying iron concentrations or pyoverdine treatments), a Kruskal–Wallis test was first applied to each metric (*r*, *K*, *t*!Z□d). Conditions yielding *p* ≤ 0.05 were considered statistically significant. Significant results were followed by a pairwise Conover–Iman post hoc test. Differences between conditions were considered significant if *p* ≤ 0.025.

## 3. Results

### 3.1. Iron availability shapes *Legionella* growth kinetics in a species-dependent manner

To determine how iron availability influences the growth dynamics of *L. pneumophila*, we monitored growth across a gradient of iron concentrations in BYEB medium (Figure 1). *L. pneumophila* exhibits a delayed lag phase in iron-depleted BYEB, but ultimately reaches comparable carrying capacity as in iron-replete conditions (Figure 1, Figure S1). In the present study, we captured the lag phase extension by calculating t_mid, which represents the time at which OD_600_ reaches half of the carrying capacity (K). In absence of significant differences in growth rate and final crop, t_mid is directly governed by differences in the lag-phase. The t_mid values of *L. pneumophila* rapidly declined with increasing iron concentration and reached a plateau at 67.1 μM iron, above which t_mid no longer decreased substantially. Growth rate (*r*) and carrying capacity (*K*) show only minor variation across iron concentrations (Figure S1), indicating that iron availability predominantly impacted the duration of the lag-phase. In particular, growth rate ranged between 0.398–0.407 h^-1^ at iron concentrations >33.5 μM, compared to cultures grown at lower iron concentration, in which the growth rate ranged between 0.376–0.385 h^-1^. All *K* values were in a similar range (OD600: 1.46-1.53), with no concentration-dependent trend.

**Figure 1.**
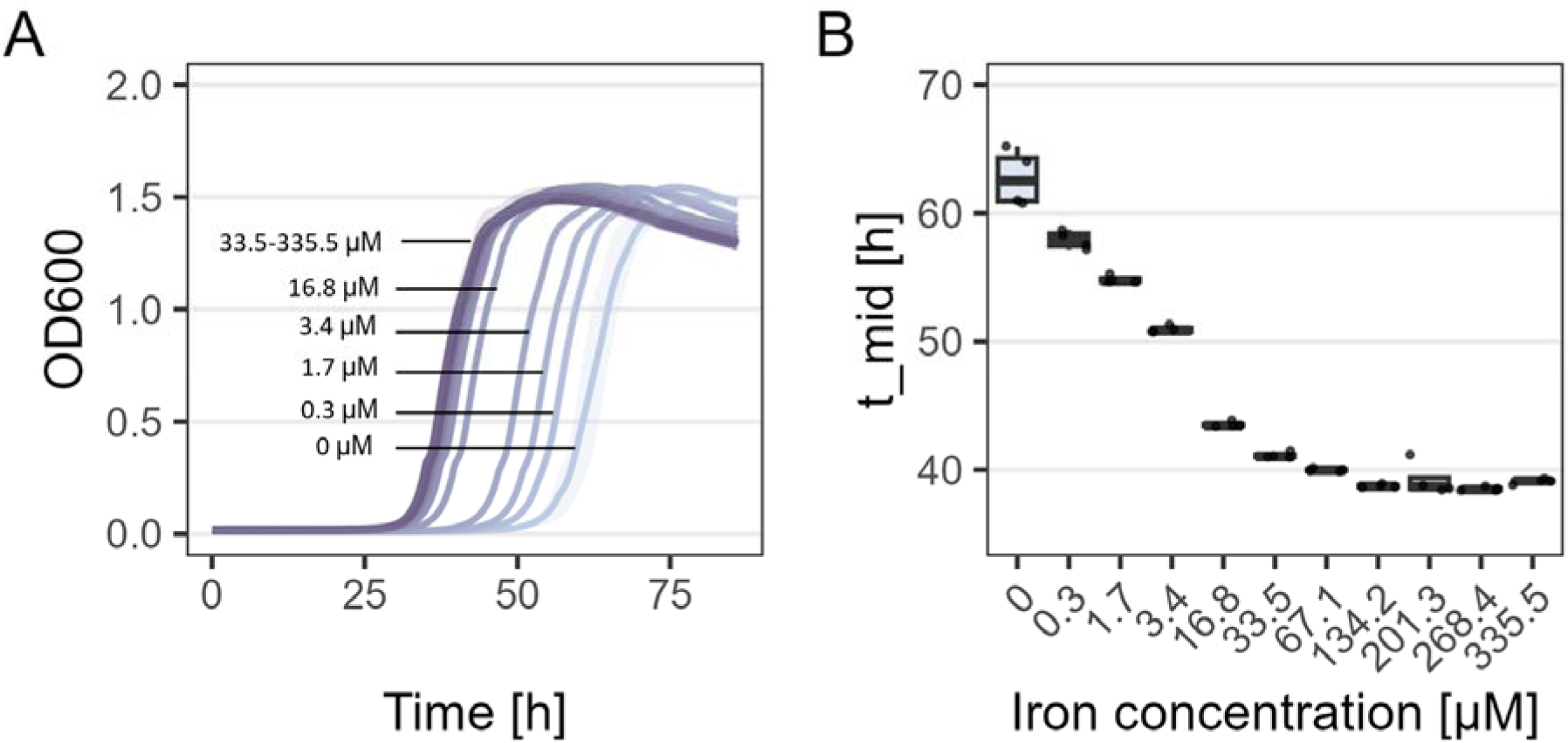
*L. pneumophila* growth in BYEB medium at different iron concentrations, recorded for 86 hours. A) The growth curves at separate iron concentrations. Each line represents the mean of 4 replicates, while the shaded areas show ±1 standard deviation from the mean. Iron concentrations [μM] are indicated next to the growth curves. B) t_mid values calculated for each iron concentration. Box plots show the first, second (median) and third quartiles, whiskers show the data range (1.5x interquartile range), dots show the individual points (n = 4).

To determine whether the observed iron-dependent growth response of *L. pneumophila* is conserved across the genus, we assessed four additional *Legionella* species under the same conditions (Figure 2, S2, S3). Iron availability strongly influenced the growth dynamics of the four *Legionella* species, with *L. feeleii* showing growth across all concentrations but at significantly higher rates (r = 0.585–0.615 h⁻¹, Figure S3) and carrying capacities only when iron concentration was above 16.8 μM. *L. jordanis* and *L. londiniensis* failed to reach stationary phase below 16.8 and 3.4 μM iron respectively, with marked improvements in growth rate and lag phase duration at higher iron concentrations (e.g., t_mid reduced from 75.5 h to 20.5 h in *L. jordanis*, p ≤ 0.025). In contrast, *L. longbeachae* maintained stable growth (r = 0.421–0.552 h⁻¹; K = 0.959–1.039) across all conditions, with only slight but significant changes in lag phase duration (max difference: 7.8 h). These inter-species differences suggest variation in iron uptake systems, storage mechanisms, or regulatory responses to iron, with implications for ecological adaptation and environmental persistence.

**Figure 2.**
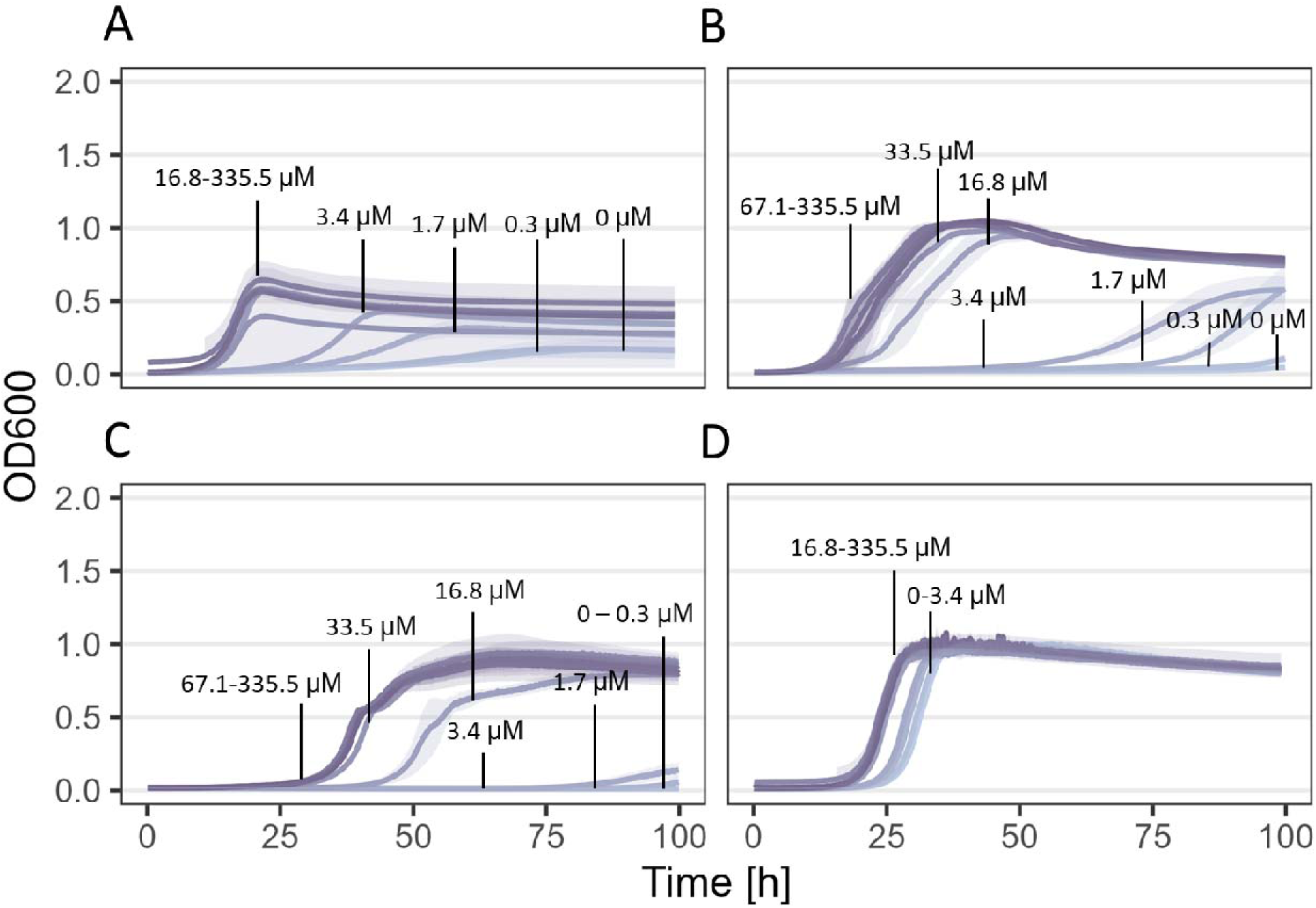
Four different *Legionella* species grown in BYEB medium at different iron concentrations, recorded for 100 hours. A) *L. feeleii*; B) *L. jordanis*; C) *L. londiniensis*; D) *L. longbeachae*. The growth curves represent the mean of 4 replicates at separate iron concentrations, while the shaded areas show ±1 standard deviation from the mean. Iron concentrations [μM] are indicated next to the growth curves.

### 3.1. Pyoverdines inhibit *L. pneumophila* growth in a concentration-dependent manner

To test whether siderophore-mediated iron sequestration limits *L. pneumophila* growth, we exposed cultures grown in BYEB medium to increasing concentrations of a commercially available mixture of pyoverdines. Indeed, we observed that t_mid values increased with rising pyoverdine concentrations (Figure 3), indicating delayed growth under stringer iron limitation. At concentrations up to 1 μM *L. pneumophila* growth was only marginally affected compared to control. In contrast, t_mid values progressively increased with higher pyoverdine concentrations (e.g., increase from 43 to 62.3 hours with 7.5 μM pyoverdine).

**Figure 3.**
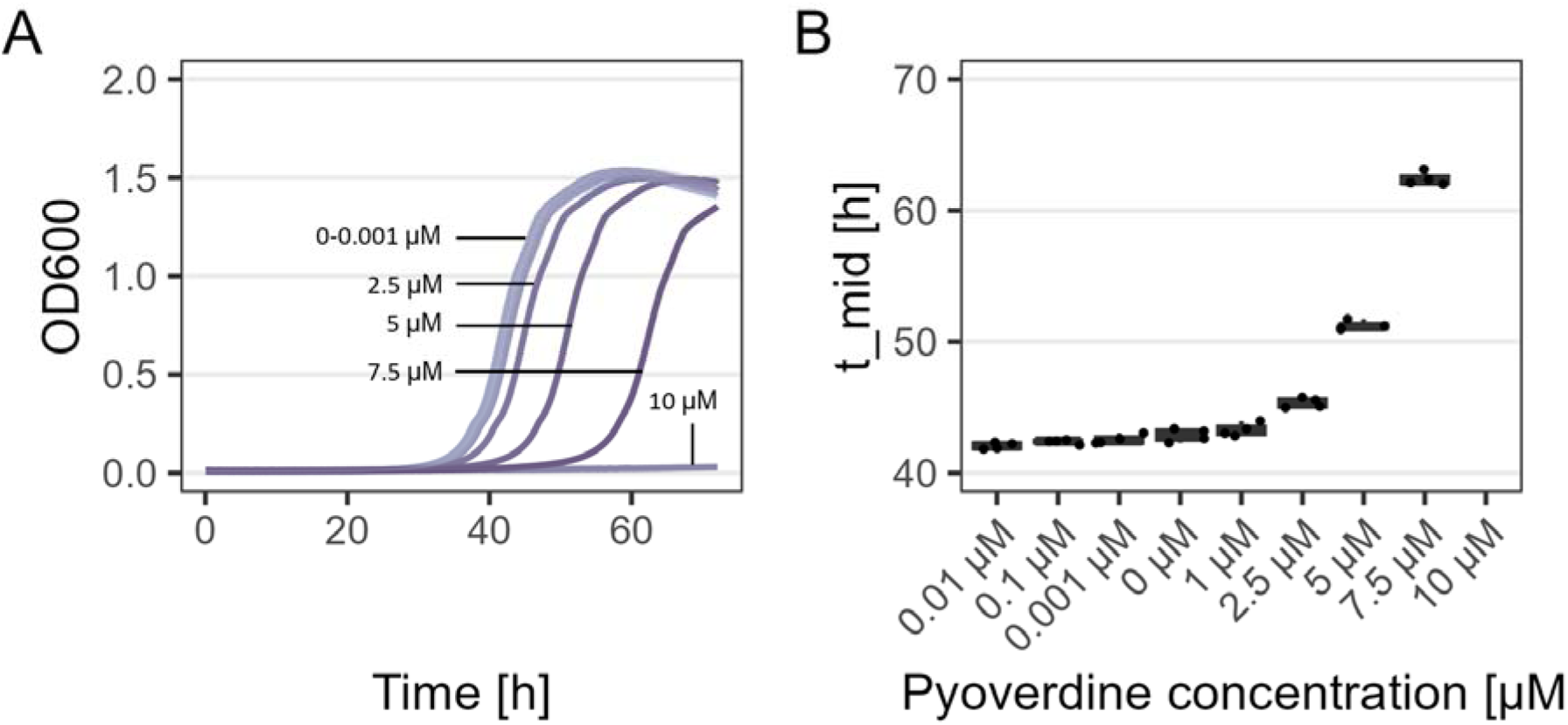
Growth of *L. pneumophila* in BYEB non-supplemented with iron at different commercial pyoverdine concentrations, recorded for 72 hours. A) Growth curves at the separate pyoverdine concentrations. A control with no pyoverdine added was included. Each line represents the mean of 3 replicates, while the shaded areas show ±1 standard deviation from the mean. Pyoverdine concentration [μg/mL] are indicated next to the growth curves. B) t_mid values calculated for each concentration. Concentrations at which *L. pneumophila* showed no growth are indicated as ND. Box plots show the first, second (median) and third quartiles, whiskers show the data range (1.5x interquartile range), dots show the raw data (n = 3).

Growth rates and K were only mildly affected up to 7.5 μg/mL (Figure S4), while we observe complete growth arrest above a threshold concentration (here 10 μg/mL). Notably, such binary response could signify that at high siderophore concentrations the iron starvation is so severe that *L. pneumophila* can no longer respond and use its own iron uptake systems. Carrying capacity trended down with increasing pyoverdine concentration: OD600 values fell from ∼1.49–1.52 at low concentrations to 1.48 at 5 μg/mL and 1.39 at 7.5 μg/mL in the presence of high pyoverdine (Figure S4). These results demonstrate that pyoverdines primarily inhibits *L. pneumophila* by extending its lag phase at low to intermediate concentrations (≤ 7.5 μg/mL), but can induce complete growth arrest at high concentrations (≥ 10 μg/mL), supporting the model that iron sequestration limits bacterial growth.

### 3.2. Supernatants from environmental *Pseudomonas* strains inhibit *L. pneumophila* growth

We then explored whether the *L. pneumophila* growth inhibition observed with the commercial pyoverdine can be reproduced with pyoverdines produced by environmental *Pseudomonas* strains from soil and freshwater habitats. In this context, it is important to note that more than 200 chemically different variants of pyoverdines exists. Here, we used five *Pseudomonas* isolates (3A06, 3G07, s3b09, s3c13, s3e20) that are known to produce a distinct pyoverdine variant each (Rehm *et al*., 2023). In a first assay, we exposed *L. pneumophila* cultured both in iron-replete and iron-depleted BYEB media to filtered supernatants harvested from these five isolates. Supernatants contain a high concentration of pyoverdines among other secreted compounds. In iron-depleted medium, four of the five *Pseudomonas* supernatants (from isolates 3A06, 3G07, s3b09, and s3e20) completely inhibited *L. pneumophila* growth (Figure 3A). Only the supernatant from s3c13 allowed limited growth, though it substantially extended the lag phase. Even in iron-supplemented BYEB, all supernatants impaired *L. pneumophila* growth compared to controls, indicating that the inhibitory effect was only partially alleviated by iron supplementation. Among all treatments, s3b09 supernatant had the strongest inhibitory effect under iron-rich conditions: growth was delayed so severely that carrying capacity was never reached within the experiment’s timeframe (72 hours). Similarly, cultures exposed to supernatants from 3A06, s3c13, s3e20, and 3G07 reached t_mid significantly later (38.0–39.6 h) than iron-supplemented controls without supernatant (t_mid = 30.1 h) or with PBS alone (31.6 h), confirming that supernatant-derived compounds are primarily responsible for the inhibitory phenotype. These results suggest that pyoverdines secreted by *Pseudomonas* strains delay or prevent *L. pneumophila* growth by effectively limiting bioavailable iron, although additional extracellular inhibitors cannot be excluded.

### 3.2. Crude pyoverdine extracts from *Pseudomonas* strains exhibit strong, strain-specific inhibition of *L. pneumophila*

To verify that pyoverdines and no other secreted compounds inhibit *L. pneumophila* growth, we crude purified the supernatants to extract pyoverdine from the five *Pseudomonas* isolates (3A06, 3G07, s3b09, s3c13, s3e20). We then tested for their inhibitory effects on *L. pneumophila* growth in both iron-limited and iron-supplemented BYEB media. We found that all crude-purified pyoverdines significantly supressed the growth of *L. pneumophila* (Figure 4). However, complete growth arrests were only observed for two pyoverdines (s3c13, s3e20), whereas a substantial extension of t_mid occurred for the remaining three (3A06, 3G07, s3b09). Similar to the supernatants, inhibition was more pronounced under iron-limited compared to iron-replete conditions, where pyoverdines more effectively scavenge available iron.

**Figure 4.**
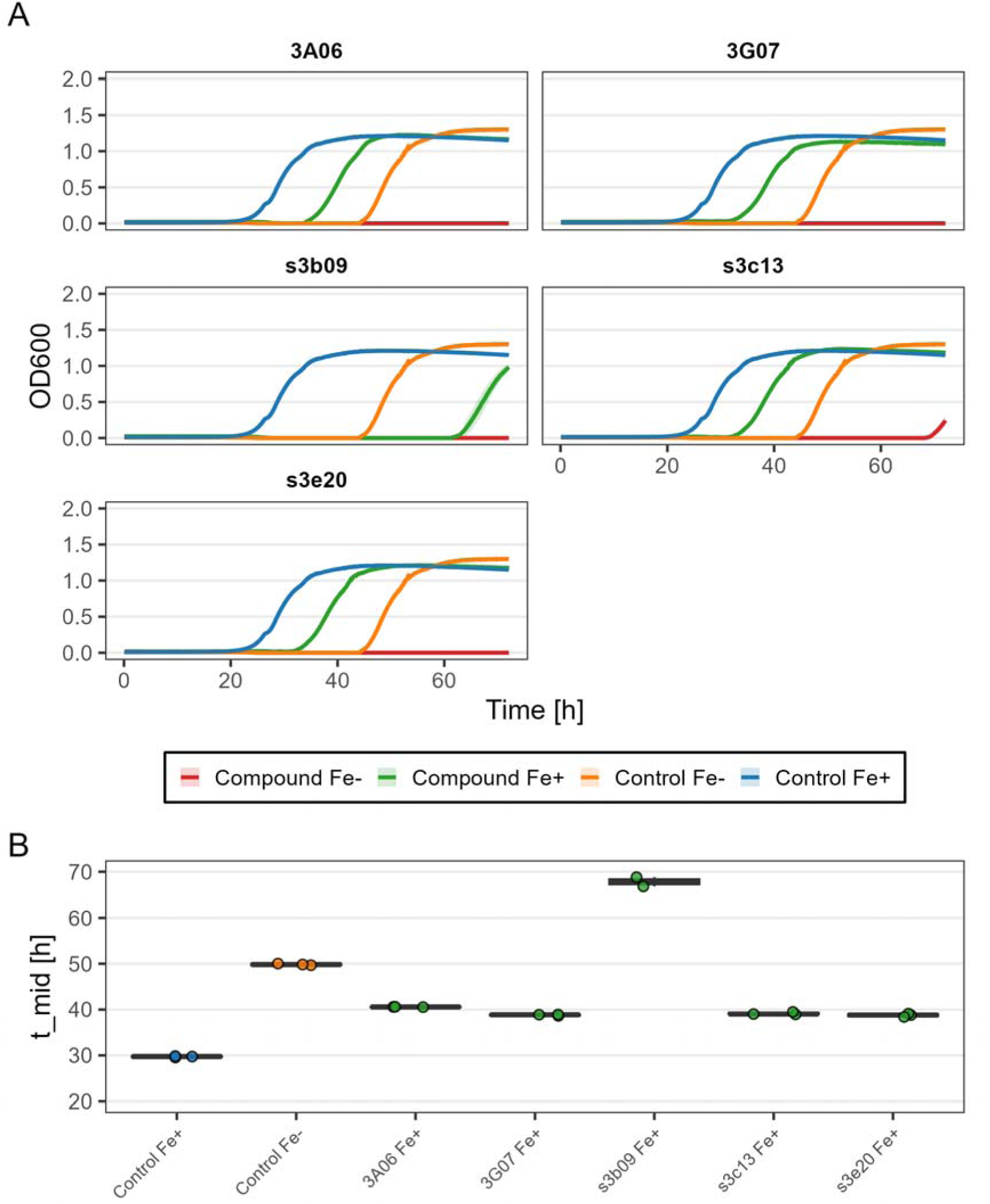
**Growth of *L. pneumophila* in supernatant of siderophores-producing *Pseudomonas***. **A)** Growth curves of *L. pneumophila* cultures grown in BYEB supplemented (compound Fe+) or unsupplemented (compound Fe-) with iron, and supernatant (SN) from five *Pseudomonas* environmental isolates, annotated on top of the respective graphs. Controls include BYEB and BYEB-Fe with no supernatant added. The individual conditions are indicated in the plot. Each line represents the mean of 3 replicates, while the shaded areas show ±1 standard deviation from the mean. **B) Box plot of** t_mid values of *L. pneumophila* cultures grown in BYEB unsupplemented and not supplemented with iron, with added supernatant from five *Pseudomonas* environmental isolates. Conditions in which no growth occurred are not displayed. Box plots show the first, second (median) and third quartiles, whiskers show the data range (1.5x interquartile range), dots show the raw data (n = 3).

**Figure 5.**
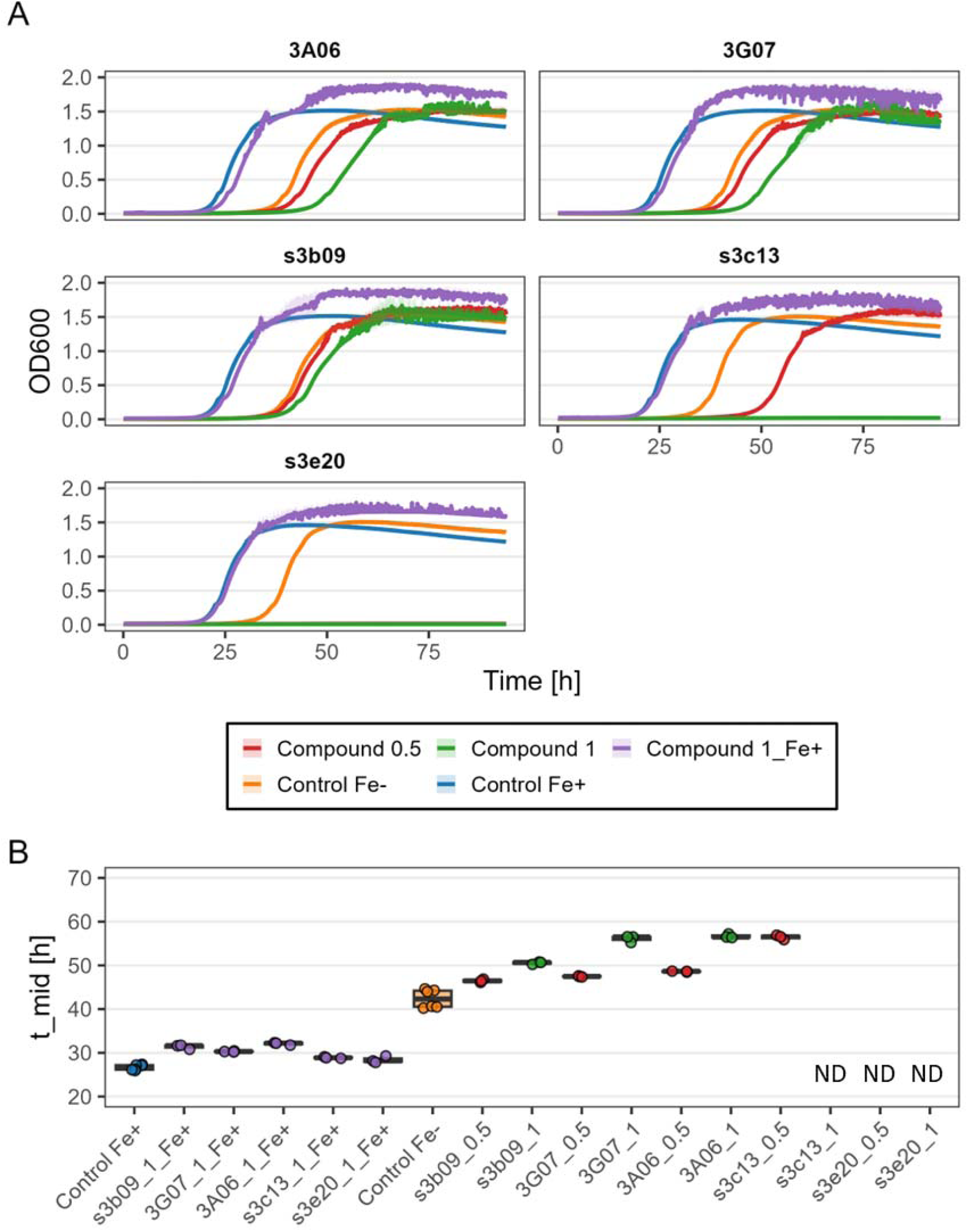
**Growth of L. pneumophila in crude-purified siderophores**. **A)** Growth curves of *L. pneumophila* cultures grown in BYEB supplemented (compound 1_Fe+) or unsupplemented (compound 0.5, compound 1) with iron, and crude purified siderophores from five *Pseudomonas* environmental isolates, annotated on top of the respective graphs. Crude purified pyoverdine were introduce at 0.5 and 1 mg/mL (compound 0.5, compound 1). Controls include BYEB and BYEB-Fe with no crude extract added. The individual conditions are indicated in the plot. Each line represents the mean of 3 replicates, while the shaded areas show ±1 standard deviation from the mean. **B) Box plot of** t_mid values of *L. pneumophila* cultures grown in BYEB unsupplemented and not supplemented with iron, with added crude purified siderophores from five *Pseudomonas* environmental isolates. Conditions in which no growth occurred are indicated with NA. Box plots show the first, second (median) and third quartiles, whiskers show the data range (1.5x interquartile range), dots show the raw data (n = 3).

The pyoverdine from the strains3b09 exhibited the weakest inhibitory effect in BYEB, with only a 5.3-hour increase in t_mid at 1 mg/mL relative to control. Pyoverdines from 3G07 and 3A06 induced stronger delays in growth onset, increasing t_mid by 10.8 and 12.1 hours, respectively, in iron-supplemented medium. The most potent inhibitory effects were observed with pyoverdines from s3c13 and s3e20: at 1 mg/mL, these extracts completely inhibited *L. pneumophila* growth. Even at a concentration of 0.5 mg/mL, s3e20 pyoverdine fully inhibited growth, while s3c13 still permitted delayed growth: in this case, t_mid was extended to 56.4 hours — 16.1 hours longer than the corresponding control. OD measurements in late exponential to early stationary phase showed fluctuations, potentially due to cell aggregation or metabolic stress, which complicated the correct estimation of growth rate and carrying capacity metrics. However, these patterns did not alter the overall trend in t_mid-based inhibition. Overall, these results show that crude pyoverdines, particularly from strains s3c13 and s3e20, are potent inhibitors of *L. pneumophila*, and more effective than filtered supernatants at equivalent concentrations.

## 4. Discussion

### 4.1. Iron availability predominantly affects the lag-phase of *Legionella* growth

Our data shows that *L. pneumophila* is capable of growing in iron-limited media. Although the lag phase is markedly extended by iron limitation, the growth rate and the carrying capacity are not. Iron has been reported to be an important nutrient for *L. pneumophila*. Early evidence clearly indicated that iron is essential for growth, with initial studies suggesting a minimal requirement as low as 3 µM for virulent strains and as high as 13 µM (Johnson *et al*., 1991, James *et al*., 1995). However, there is no clear consensus on the precise iron requirement, with reports ranging broadly from about 1 µM up to over 20 µM (Reeves *et al*., 1981), depending on experimental conditions and strain-specific virulence. Moreover, it has been demonstrated that iron is crucial for the intracellular replication of *Legionella*, with dedicated effectors that promote the delivery of iron to the *Legionella*-containing-vacuole (Isaac *et al*., 2015) and low intracellular iron levels triggering the escape of the *Legionella* from its host (O’Connor Tamara *et al*., 2016). While the exact role of iron in *Legionella* biology has to date not been evaluated, it has been documented that for most bacteria, iron is present in respiratory and metabolic proteins as part of Fe-S clusters, which are essential for the action of specific enzymes (Rolfe Matthew *et al*., 2012). Consistently, it has been described that in *L. pneumophila* the protein with the highest iron content is in fact an aconitase (Mengaud & Horwitz, 1993).

In our experiments, we showed that decreasing the iron concentration in the media caused a progressive extension of the lag phase. This is consistent with previous work by Rolfe and colleagues, who observed that bacteria accumulate iron during the lag phase in order to prepare metabolic proteins for the exponential phase (Rolfe Matthew *et al*., 2012). In particular, the authors demonstrated that such process is accompanied by the upregulation of genes related to expression of iron transporters and siderophores (Rolfe Matthew *et al*., 2012). We demonstrated that upon acquisition of the iron needed, the growth proceeds without impacting the final yield. In fact, our experiments showed that *Legionella* harvested in exponential phase did have an extended lag-phase when re-inoculated in iron-depleted conditions (Figure S5). At the same time, evidence shows that the host can limit iron using e.g., transferrin as an immune system against infection. In this context, iron accumulation during the lag phase would represent a counter strategy to the defense mechanisms exhibited by the host. Interestingly, this process has been shown primarily for pathogenic bacteria (Bertrand, 2014). We furthermore explored for the first time the iron response of different *Legionella* species. Across tested *Legionella* species, however, responses to iron supplementation were not uniform: some species showed modest changes in growth dynamics, while others demonstrated stronger dependence on iron for growth rate or yield. While it is known that differences are present among species of *Legionella* with respect to iron acquisition strategies, it is not clear whether iron is used differently across the genus. However, it is safe to assume that these inter-species differences likely reflect variation in ecological niches, or degrees of adaptation to intracellular versus extracellular lifestyles. In the case of iron, given its probable involvement in metabolic processes, it is possible to argue that *Legionella* species are metabolically different, as already demonstrated by our group in a deep pangenomic analysis of the *Legionellaceae* family (Gabrielli *et al*., 2024).

### 4.1. Pyoverdines delay *L. pneumophila* growth

The addition of commercial pyoverdines resulted in dose-dependent inhibition of *L. pneumophila* growth, with a clear effect on t_mid (lag phase extension) at concentrations ≥1 μg/mL. Moreover, supernatants and crude extracts from pyoverdine-producing environmental *Pseudomonas* strains inhibited the growth of *L. pneumophila*, particularly in iron-limited conditions, where complete inhibition occurred in most treatments. Notably, the magnitude of inhibition was strain-specific, consistent with differences in the chemical structure of the pyoverdine variants produced by each strain (Vollenweider *et al*., 2024). This adds to previous observations of *Pseudomonas* producing multiple siderophores and switch among them to increase the chance of outcompeting bacteria with similar siderophores or less potent ones (Leinweber *et al*., 2018), which gives them a strong advantage in respects to iron competition. It is important to note that, while the supernatant experiments might have been influenced by other inhibitory compounds produced by *Pseudomonas* spp. (Cavallaro *et al*., 2025), this is not the case in case of the addition of crude-purified pyoverdine, confirming the inhibitory action by iron deprivation.

*L. pneumophila* has also been found to produce at least one siderophore, namely Legiobactin, with identical chemical structure to Rhizoferrin, a siderophore produced by several fungi and bacteria (Liles *et al*., 2000, Burnside *et al*., 2015). This would, in theory, enable competition with *Pseudomonas* siderophores for available iron. However, it is notable that different siderophores have specific potency, which can be indicated as their stability constant K. While no information on the stability constant of legiobactin is to date available, it is known that Rhizoferrin has a K = 19 (Yehuda *et al*., 2000). Pyoverdine is reported to have a stability constant K = 32 (Meyer & Abdallah, 1978, Vollenweider *et al*., 2024), it is hence safe to assume that pyoverdine-producing *Pseudomonas* would be able to outcompete *Legionella* through the production of a more potent siderophore. Previous studies have already demonstrated that siderophores produced by different bacterial species can act as inhibitors the growth of other pathogenic strains. For example, early investigations showed that strains of *Pseudomonas* were able to produce growth-promoting effects by inhibiting plant pathogens through iron sequestration using the siderophore pseudobactin (Kloepper *et al*., 1980). Gokarn and Pal (Gokarn & Pal, 2018) demonstrated that the siderophores exochelin-MS and deferoxamine-B, isolated from *Mycobacterium smegmatis* and *Streptomyces pilosus* respectively, were able to inhibit several strains of methicillin resistant *Staphylococcus aureus* (MRSA) and a broad range of metallo-beta-lactamase-producing *P. aeruginosa* and *Acinetobacter baumannii*, either alone or in combination with antibiotics. Moreover, growth experiments in the presence of siderophores showed, in some instances, a delay in the lag phase consistent with our observations. An additional study described the inhibitory properties of enterobactin and salmochelin S4, siderophores produced by some *Enterobacteriaceae*, towards *S. aureus* (Davidov *et al*., 2025). Specifically, the two siderophores showed inhibition with IC50 values ranging 2-5 μg/mL (salmochelin) and 5-10 μg/mL (enterobactin), indicating high potency at low concentrations. A combination of the two siderophores led to increased inhibition of the bacteria tested. More notably, the recent work of Vollenweider and coworkers investigated the ability of several *Pseudomonas* isolates (including the same strains used in this study) to inhibit a panel of human opportunistic pathogenic bacteria, including *A. baumannii, Klebsiella pneumoniae, S. aureus* and *Burkholderia cenocepacia* (Vollenweider *et al*., 2024). The authors were able to demonstrate the inhibition of 12 opportunistic pathogenic species with 25 siderophores-producing *Pseudomonas*, with the growth restored after the addiction of FeCl₃, confirming iron dependency. Purified pyoverdines completely inhibited the growth of *S. aureus* and *A. baumannii*, while also increasing survival in larvae infected with the pathogens and showing low toxicity towards mammalian cell lines. Interestingly, while bacteria naturally produce siderophores in iron-depleted conditions, the isolated siderophores tested in our work produced an effect on bacterial growth even in iron-replete conditions. In fact, our experiments using the supernatant and the crude extract from the *Pseudomonas* strains, showed an extension of the lag-phase even when iron is added to the medium containing the siderophores.

### 4.1. Implications for controlling *Legionella* in built environments

Our results support the concept that siderophore-producing bacteria, or purified siderophores themselves, could serve as biological antagonists to *Legionella* in the environment, especially in the extracellular phase of their biphasic lifecycle (Barbosa *et al*., 2024), when the protist would not be able to provide iron intracellularly. In previous work (Cavallaro *et al*., 2022), we have reviewed the underlying mechanisms that may be responsible for the natural inhibition of *Legionella* by some bacterial taxa and microbial consortia. There, given the well-known iron dependencies of *Legionella*, we hypothesized that siderophore-producing bacteria could potentially be suitable for a probiotic approach, however, no data virtually existed at the time. The findings presented in the present work reinforce this hypothesis and point towards exploitative competition for specific resources being an important tool for *Legionella* control. However, many antagonism studies are focused on interference competition, which often offer a simpler experimental design and more straightforward results. In this regard, siderophores represent a nice opportunity, because the exploitative competition is performed by means of a secreted compound; this offers the possibility to isolate the siderophores, conduct dedicated tests, and design deliverable control strategies. From an applicative perspective, this could form the basis for passive control strategies in water systems, for example, bioaugmentation with siderophore-producers or disinfection-based methods that use siderophores to scavenge iron from the environment and limit the occurrence of problematic organisms. This approach has already been investigated for diverse applications. For example, Kamath and colleagues (Kamath *et al*., 2025) showed that purified siderophores from *Bacillus amyloliquefaciens* D5 reduced leaf spot severity in mung bean by depriving *Cercospora canescens* of iron, which coincided with increased host defense enzyme activity. Dimopoulou et al. (Dimopoulou *et al*., 2021) further demonstrated that bacillibactin from *B. amyloliquefaciens* MBI600 directly inhibited *Pseudomonas syringae* in tomato, underscoring how siderophore-mediated iron competition can suppress bacterial pathogens in planta. Siderophore-producing strains have also proven effective in diverse environments: *Trichoderma virens* XZ11-1 lowered Fusarium wilt incidence in banana while promoting plant growth (Cui *et al*., 2025), a marine *Glutamicibacter* sp. reduced mortality from shrimp-pathogenic vibrios in aquaculture through iron sequestration (Shi *et al*., 2025), and *Bifidobacterium* strains with high iron-binding capacity limited the growth of enteric pathogens in gut cell models (Vazquez-Gutierrez *et al*., 2016). Beyond pathogen control, siderophores are also being explored for bioremediation, as they can chelate a range of metals and thus contribute to pollutant detoxification and enhanced plant metal tolerance (Kurth *et al*., 2016, Wei *et al*., 2025). Importantly, this mechanism circumvents the need for bactericidal action and would shift instead the competitive behaviour in a way that disfavours *Legionella* and is less disruptive of the surrounding microbiota, although still untargeted in its mode of action. Finally, in engineered aquatic ecosystems, where iron can leach by some of the pipe materials (e.g., following corrosion) and hence be present in higher concentrations in the water, this might lead to increased growth of *Legionella*. Hence, control strategies through siderophores could represent a valuable strategy to limit the amount of iron in the water and avoid *Legionella* proliferation.

## Conclusions

- *L. pneumophila* growth was susceptible to low iron concentrations, with the lag phase being the main parameter affected.
- Different species of *Legionella*, however, show a diverse response, which might be linked to differences in iron utilization.
- The siderophore pyoverdine was able to inhibit the growth of *L. pneumophila* in iron limited environments, where a primary effect was observed on the lag-phase. At high concentrations of pyoverdine, the growth of *L. pneumophila* was completely inhibited.
- Extracellular extracts and crude-purified pyoverdines from environmental *Pseudomonas* were able to delay the lag phase of *L. pneumophila* in iron repleted conditions and to completely inhibit its growth at low iron concentrations. This suggests that in the environment, siderophores producing bacteria might outcompete *Legionella* for iron acquisition.
- While the production of siderophores is generally driven by iron scarcity, purified siderophores might be used as an alternative mitigation strategy against *Legionella*, by scavenging environmental iron in engineered aquatic ecosystems.

