## Supplementary Information for "Siderophore-mediated inhibition of *Legionella pneumophila* by environmental *Pseudomonas* isolates"

* Corresponding author:

Name:	Frederik Hammes

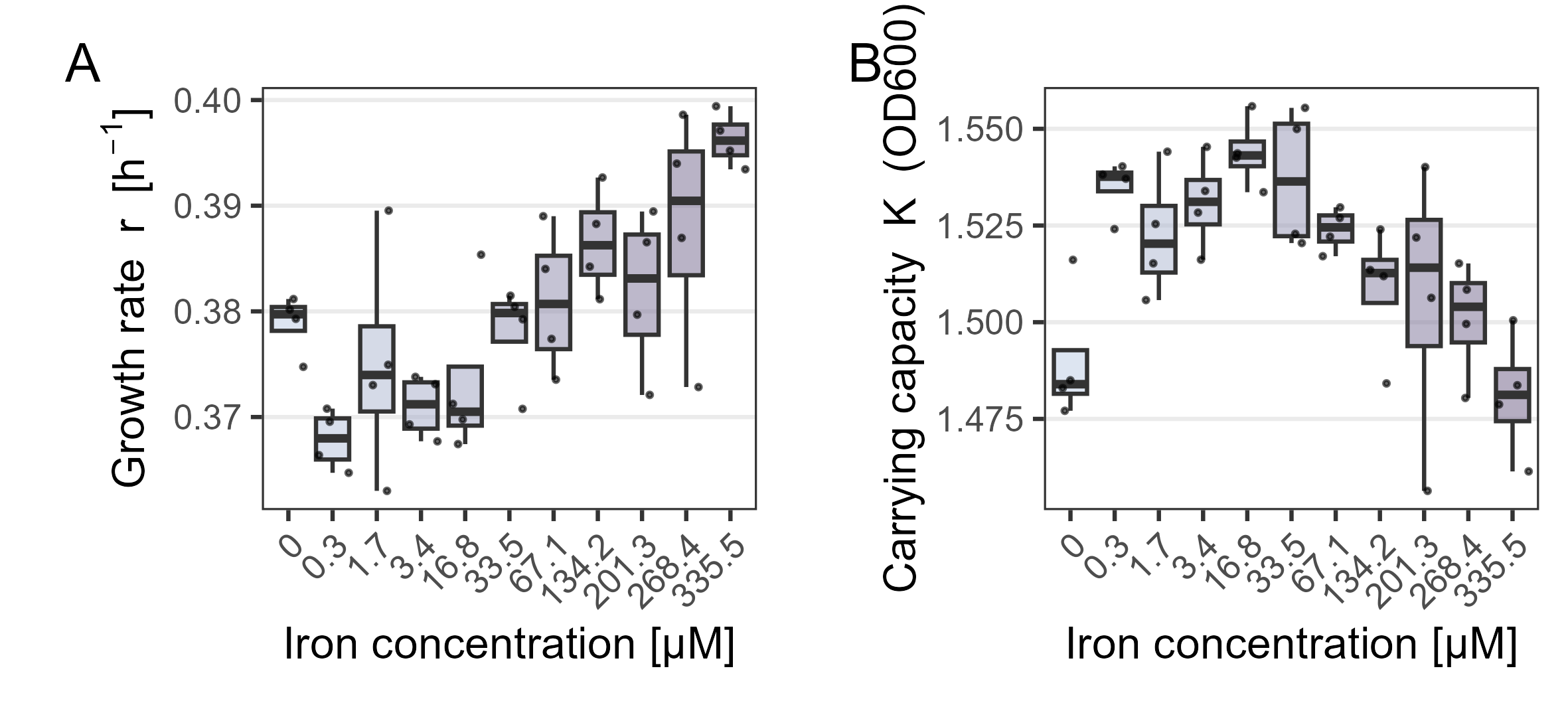

### **Figure S1. *L. pneumophila* growth parameters in BYEB medium at different iron concentrations.**

(A) Growth rate (r, h⁻¹) and (B) carrying capacity (K, OD600) estimated from OD600 time-series data at increasing iron concentrations. Iron concentrations [µM] are shown on the x-axis. Growth parameters were obtained by fitting logistic growth models to replicate-level growth curves. Box plots show the first, second (median), and third quartiles; whiskers indicate the data range (1.5× interquartile range), and dots represent individual replicate fits.

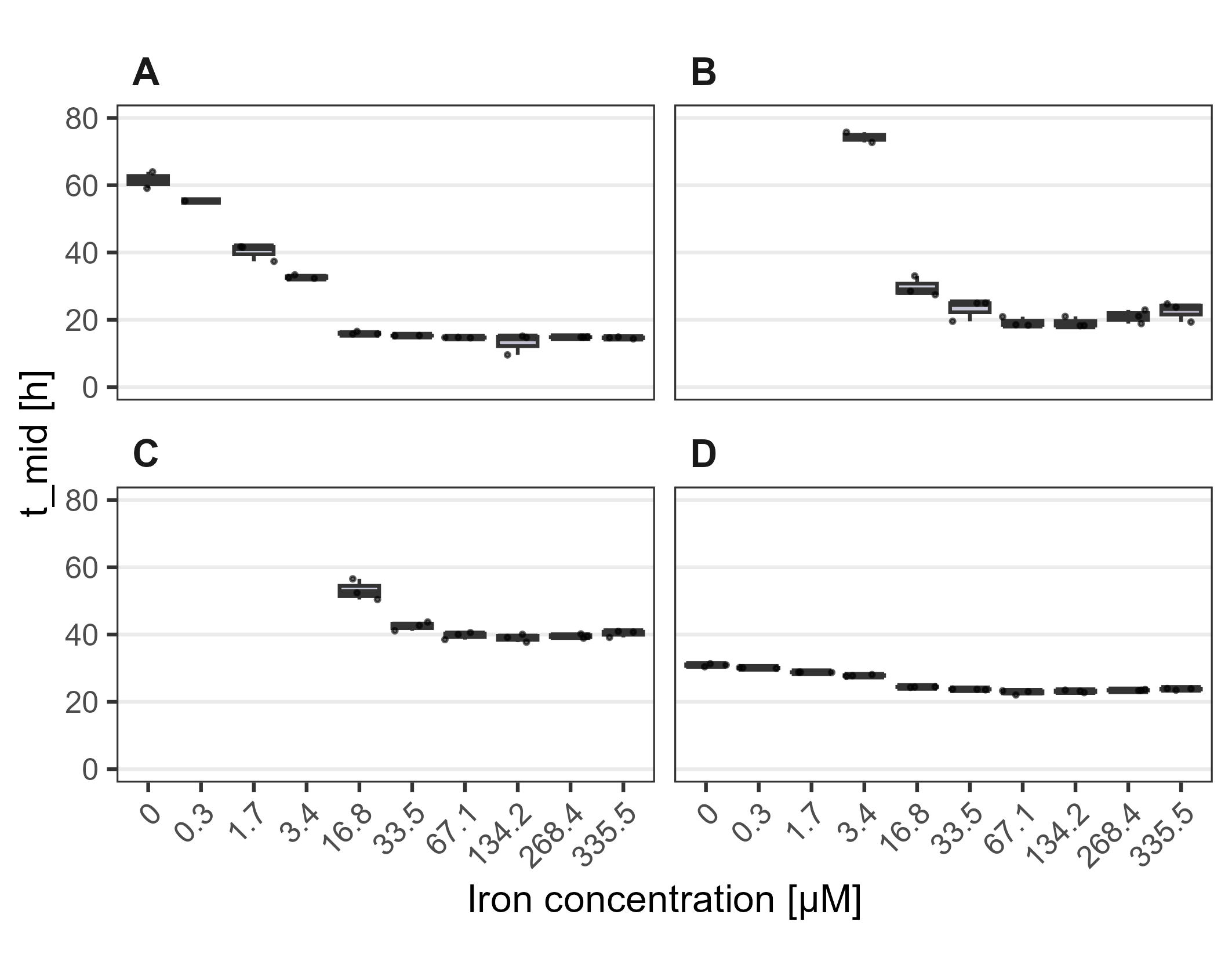

ND

ND

ND

ND

ND

ND

### **Figure S2. Species-specific t_mid values for four *Legionella* species across iron concentrations.**

*t*_mid values were calculated for *Legionella* strains grown in BYEB medium at increasing iron concentrations. Iron concentrations [µM] are shown on the x-axis. Panels correspond to individual species as follows: panel A, *L. feeleii*; panel B, *L. jordanis*; panel C, *L. londiniensis*; and panel D, *L. longbeachae*. Box plots show the first, second (median), and third quartiles; whiskers indicate the data range (1.5× interquartile range), and dots represent individual replicate-level model fits (n = 3). *t*ₘᵢd values were derived from logistic growth models fitted to replicate-level OD600 time-series data. Replicates were excluded from the analysis when no detectable growth was observed, defined as fits yielding a carrying capacity (*K*) below 0.2 OD600.

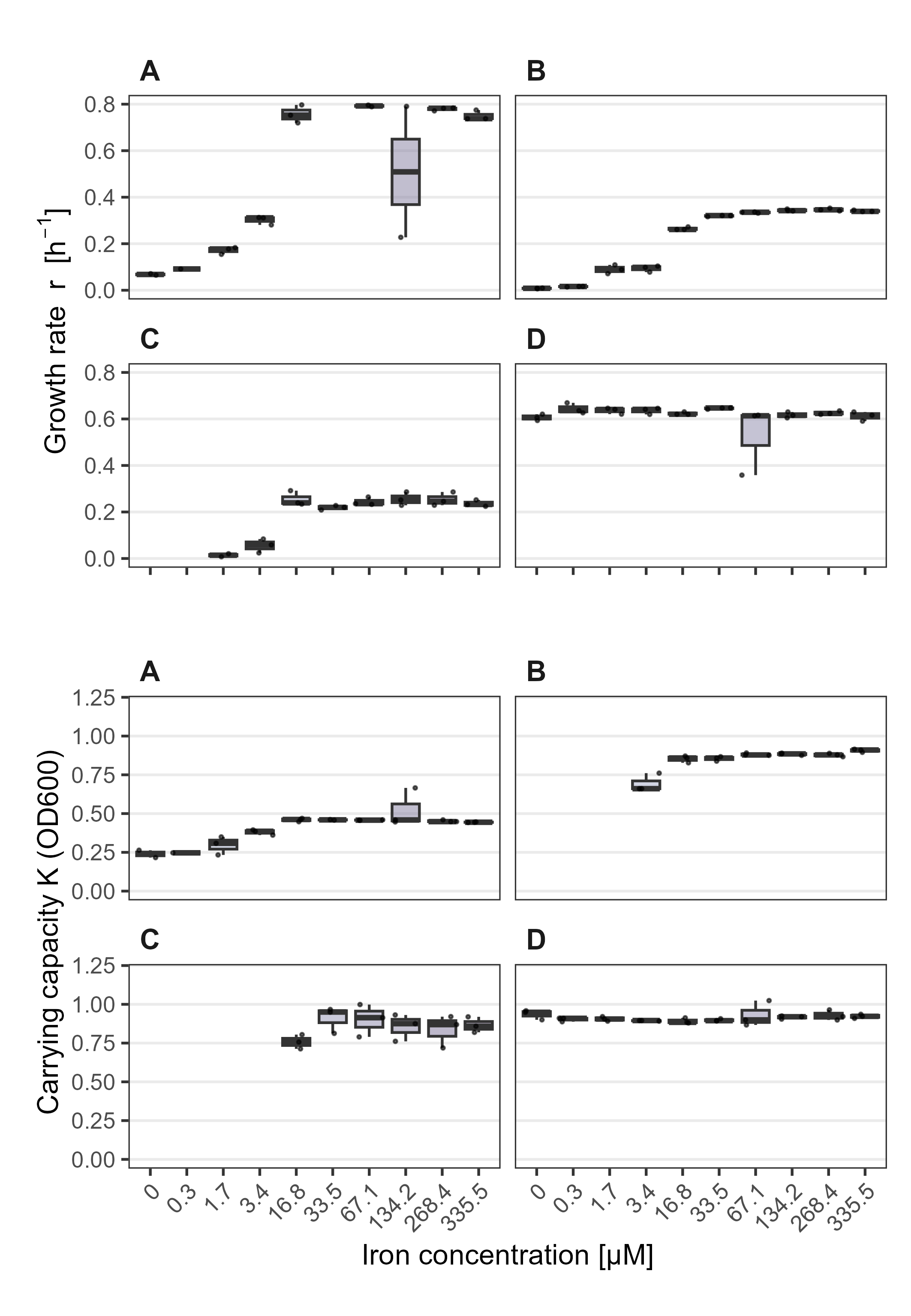
**Figure S3. Species-specific growth rate (*r*) and carrying capacity (*K*) for four *Legionella* species across iron concentrations.**

ND

ND

ND

ND

ND

ND

ND

ND

Growth rate (*r*, h⁻¹; panel A) and carrying capacity (*K*, OD600; panel B) were calculated for *L. feeleii*, *L. jordanis*, *L. londiniensis*, and *L. longbeachae* grown in BYEB medium at increasing iron concentrations. Iron concentrations [µM] are shown on the x-axis. In both panels, subpanels correspond to individual species as follows: panel A, *L. feeleii*; panel B, *L. jordanis*; panel C, *L. londiniensis*; and panel D, *L. longbeachae*. Growth parameters were obtained by fitting logistic growth models to replicate-level OD600 time-series data. Box plots show the first, second (median), and third quartiles; whiskers indicate the data range (1.5× interquartile range), and dots represent individual replicate-level model fits (n = 3). Replicates without detectable growth were excluded, defined as logistic fits yielding a carrying capacity (*K*) below 0.2 OD600; additionally, unrealistically large *K* estimates (> 2 OD600), indicative of non-identifiable model fits under weak-growth conditions, were excluded from the analysis.

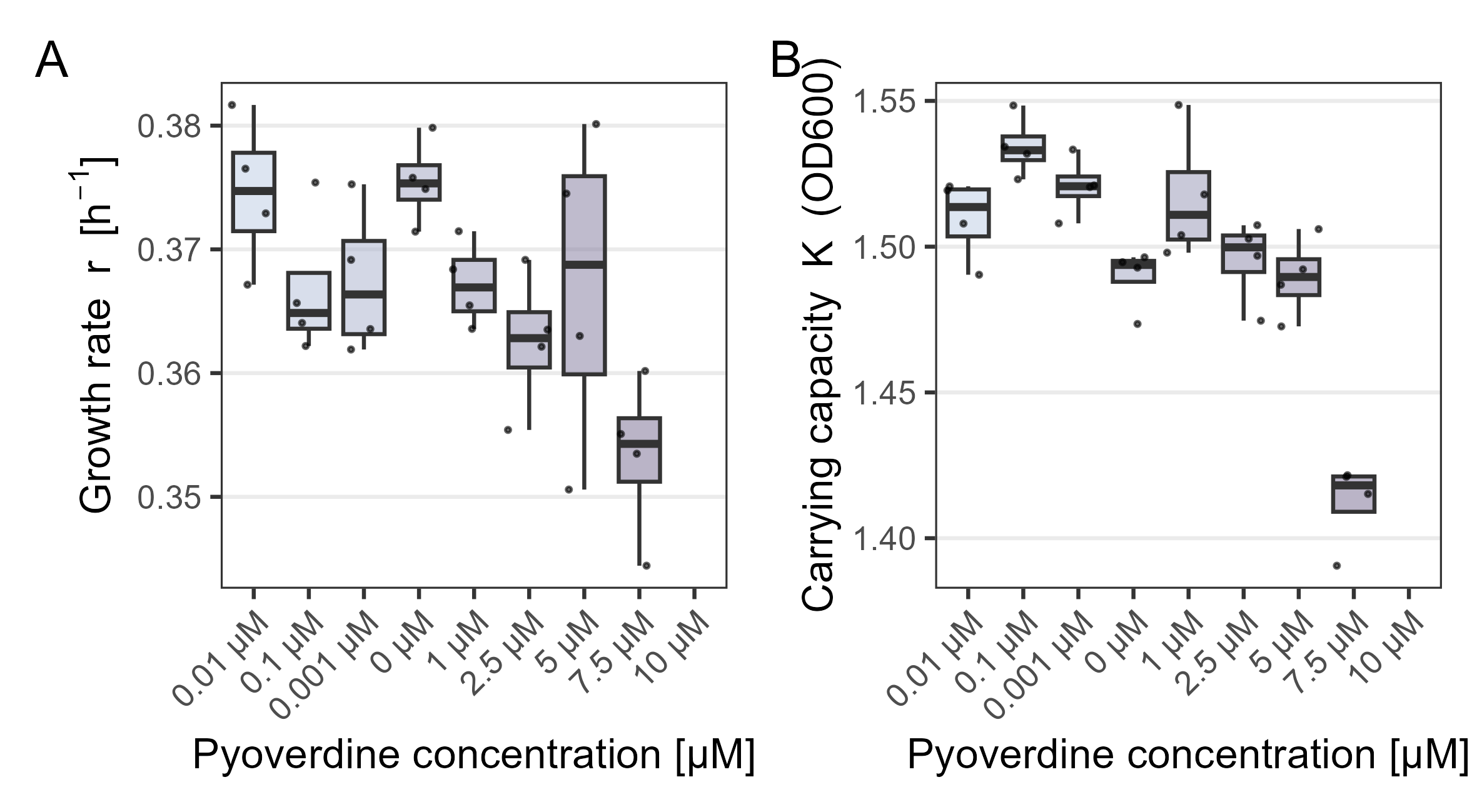

### **Figure S4. Effect of commercial pyoverdine on *L. pneumophila* growth parameters.**

Growth rate (*r*, h⁻¹; panel A) and carrying capacity (*K*, OD600; panel B) were determined for *L. pneumophila* grown in BYEB medium in the presence of increasing concentrations of commercial pyoverdine. Growth parameters were obtained by fitting logistic growth models to OD600 time-series data. Box plots show the first, second (median), and third quartiles; whiskers indicate the data range (1.5× interquartile range), and dots represent individual replicate-level model fits (n = 4). Replicates without detectable growth were excluded from the analysis, defined as fits yielding a carrying capacity (*K*) below 0.2 OD600.

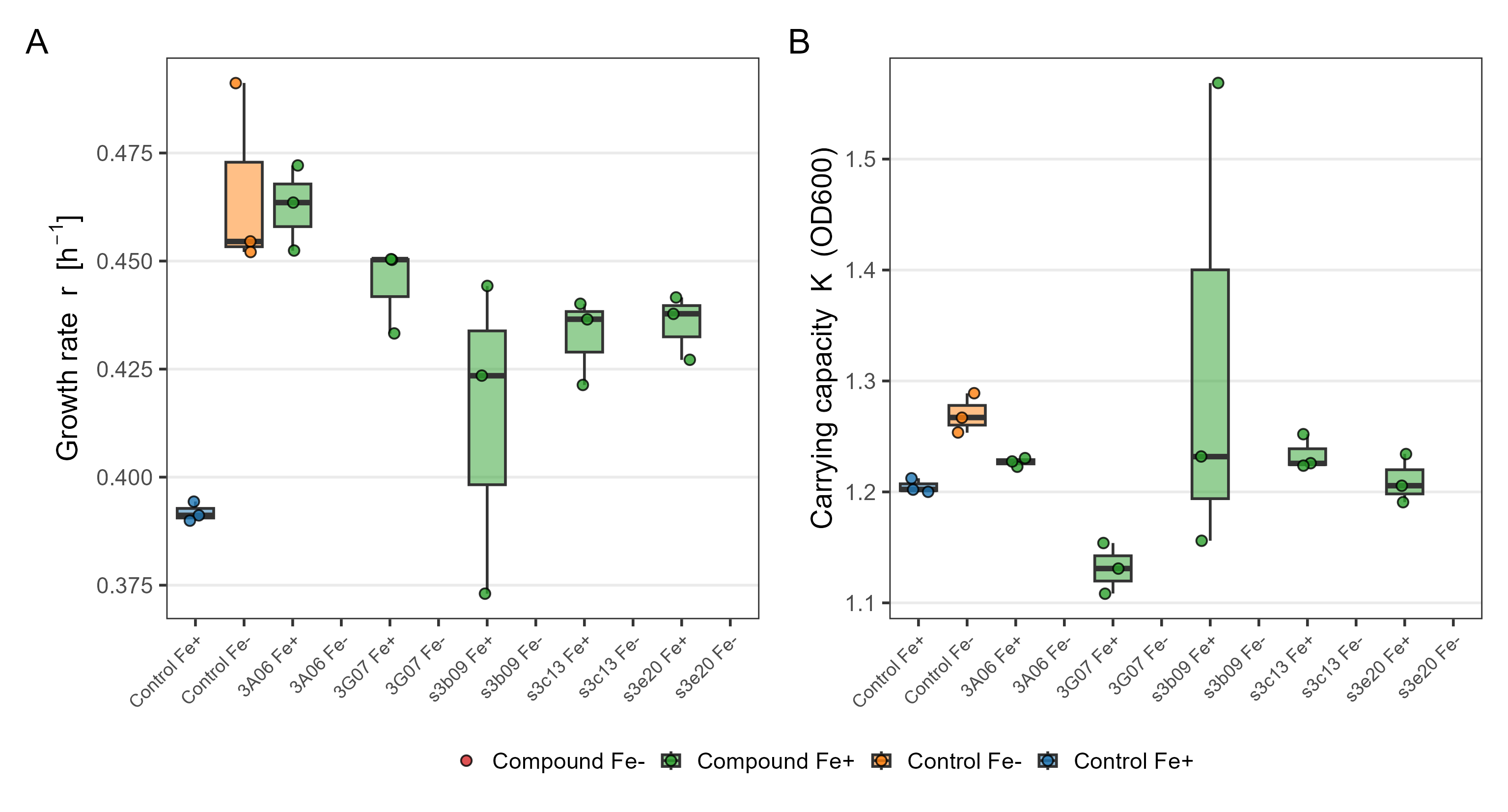

ND

ND

ND

ND

ND

ND

ND

ND

ND

ND

### **Figure S5. Growth rate (*r*) and carrying capacity (*K*) of *L. pneumophila* in supernatants from siderophore-producing *Pseudomonas* isolates under iron-replete and iron-depleted conditions.**

Growth rate (*r*, h⁻¹; panel A) and carrying capacity (*K*, OD600; panel B) were determined for *L. pneumophila* cultures grown in BYEB medium supplemented with supernatant from five environmental *Pseudomonas* isolates, under iron-replete (Fe⁺) or iron-depleted (Fe⁻) conditions. Control conditions correspond to *L. pneumophila* grown in BYEB medium with iron (Control Fe⁺) or without iron (Control Fe⁻), in the absence of supernatant. Individual conditions, including supernatant source and iron availability, are indicated on the x-axis. Growth parameters were obtained by fitting logistic growth models to replicate-level OD600 time-series data. Box plots show the first, second (median), and third quartiles; whiskers indicate the data range (1.5× interquartile range), and dots represent individual replicate-level model fits (n = 4). Conditions in which no detectable growth occurred were excluded from the analysis, defined as fits yielding a carrying capacity (*K*) below 0.2 OD600

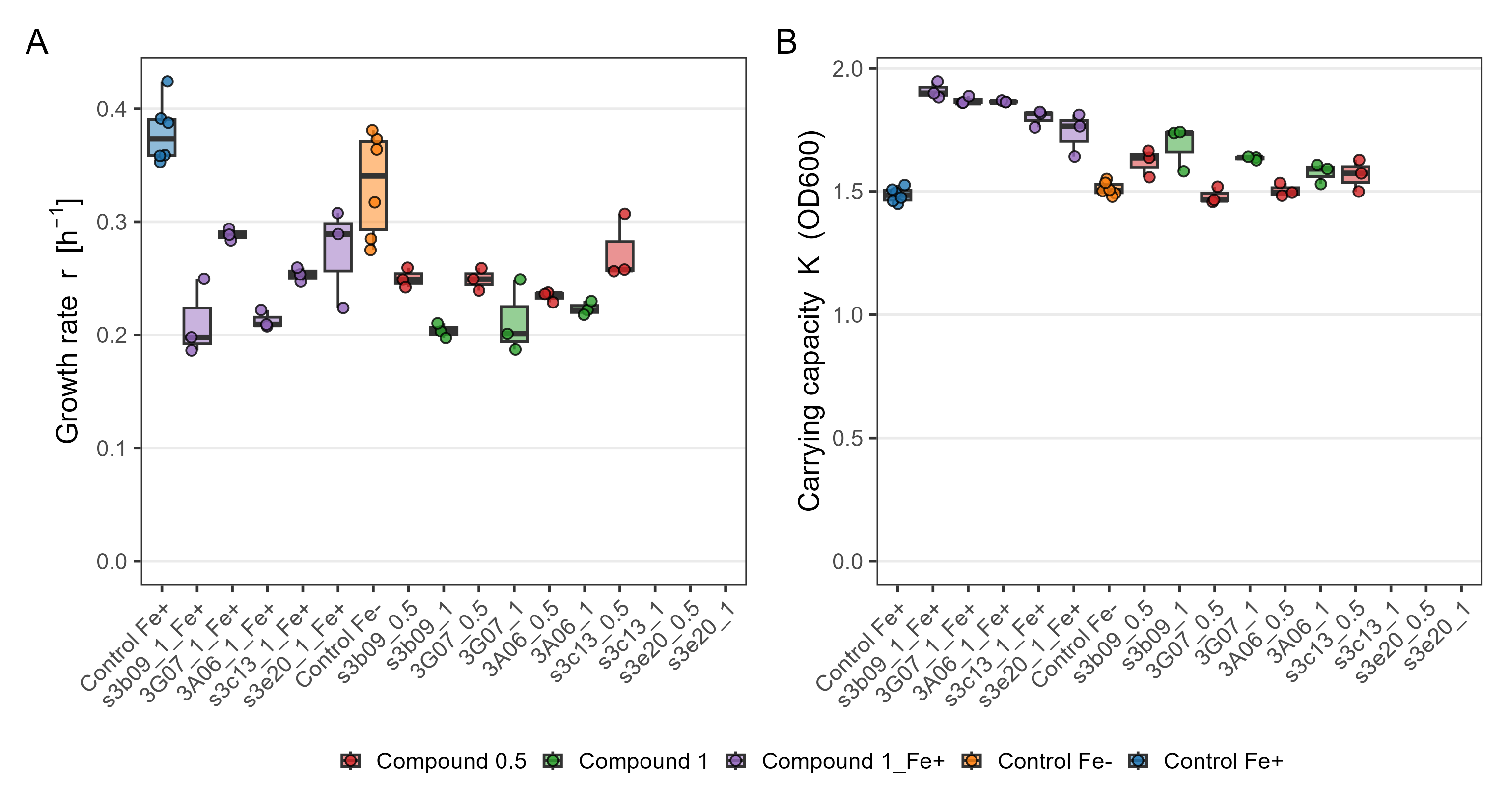

ND

ND

ND

ND

ND

### **S6. Growth rate (*r*) and carrying capacity (*K*) of *L. pneumophila* in the presence of crude-purified siderophores from *Pseudomonas* isolates under different iron conditions.**

Growth rate (*r*, h⁻¹; panel A) and carrying capacity (*K*, OD600; panel B) were determined for *L. pneumophila* cultures grown in BYEB medium supplemented with crude-purified siderophores from five environmental *Pseudomonas* isolates. Crude-purified siderophores were added at 0.5 mg mL⁻¹ (compound 0.5) or 1 mg mL⁻¹ (compound 1) under iron-depleted conditions, or at 1 mg mL⁻¹ under iron-replete conditions (compound 1_Fe⁺). Control conditions correspond to *L. pneumophila* grown in BYEB medium with iron (Control Fe⁺) or without iron (Control Fe⁻), in the absence of crude-purified siderophores. Individual conditions are indicated on the x-axis. Growth parameters were obtained by fitting logistic growth models to replicate-level OD600 time-series data. Box plots show the first, second (median), and third quartiles; whiskers indicate the data range (1.5× interquartile range), and dots represent individual replicate-level model fits (n = 3). Replicates without detectable growth were excluded from the analysis, defined as fits yielding a carrying capacity (*K*) below 0.2 OD600.
